# Sociodemographic patterning in the oral microbiome of a diverse sample of New Yorkers

**DOI:** 10.1101/189225

**Authors:** Audrey Renson, Heidi E. Jones, Francesco Beghini, Nicola Segata, Christine P. Zolnik, Mykhaylo Usyk, Thomas U. Moody, Lorna Thorpe, Robert Burk, Levi Waldron, Jennifer B. Dowd

## Abstract

1

**Purpose:** Variations in the oral microbiome are potentially implicated in social inequalities in oral disease, cancers, and metabolic disease. We describe sociodemographic variation of oral microbiomes in a diverse sample.

**Methods:** We performed 16S rRNA sequencing on mouthwash specimens in a subsample (n=282) of the 2013-14 population-based New York City Health and Nutrition Examination Study (NYC-HANES). We examined differential abundance of 216 operational taxonomic units (OTUs), and alpha and beta diversity by age, sex, income, education, nativity, and race/ethnicity. For comparison, we also examined differential abundance by diet, smoking status, and oral health behaviors.

**Results:** 69 OTUs were differentially abundant by any sociodemographic variable (false discovery rate < 0.01), including 27 by race/ethnicity, 21 by family income, 19 by education, three by sex. We also found 49 differentially abundant by smoking status, 23 by diet, 12 by oral health behaviors. Genera differing for multiple sociodemographic characteristics included *Lactobacillus*, *Prevotella*, *Porphyromonas*, *Fusobacterium*.

**Conclusions:** We identified oral microbiome variation consistent with health inequalities, with more taxa differing by race/ethnicity than diet, and more by SES variables than oral health behaviors. Investigation is warranted into possible mediating effects of the oral microbiome in social disparities in oral, metabolic and cancers.

**Highlights:**

- Most microbiome studies to date have had minimal sociodemographic variability, limiting what is known about associations of social factors and the microbiome.
- We examined the oral microbiome in a population-based sample of New Yorkers with wide sociodemographic variation.
- Numerous taxa were differentially abundant by race/ethnicity, income, education, marital status, and nativity.
- Frequently differentially abundant taxa include *Porphyromonas*, *Fusobacterium*, *Streptococcus*, and *Prevotella*, which are associated with oral and systemic disease.
- Mediation of health disparities by microbial factors may represent an important intervention site to reduce health disparities, and should be explored in prospective studies.

## 2 Introduction

Health disparities by race/ethnicity, socioeconomic status (SES), sex, and other sociodemographic factors have long been observed but their mechanisms have yet to be fully elucidated. In particular, racial/ethnic and socioeconomic disparities have been consistently observed in oral health outcomes (1), cardiovascular disease (CVD) (2, 3), diabetes (4), preterm birth and low birth weight (5, 6), and rheumatoid arthritis (7).

Variations in human oral microbiome structure and function have been associated with oral disease (8, 9), as well as a wide range of systemic illnesses including CVD (10-12), diabetes (13, 14), cancers (15-18), birth outcomes (19, 20), and rheumatoid arthritis (21, 22). Hypothesized pathways for such associations include both direct virulence and modulation of systemic immune response (15), although causal evidence is limited. Also, regardless of their causal role, the microbiota represent potentially useful biomarkers for early disease detection and risk prediction.

This combination of findings has led researchers to call for investigation into the role of the microbiome in health disparities (23) but little empirical work has yet been done in this area. A number of mechanisms potentially link social inequality to the microbiome (24). Mechanisms linking the social environment to microbe exposure have been discussed in relation to common pathogens such as CMV and EBV; these may include household crowding, use of public transportation, and differences in susceptibility due to e.g. breastfeeding (antibodies) and poor sleep (25, 26), mechanisms which may apply to commensal microbes as well. Changes in immune function related to psychosocial stress (27), nutrition (28), smoking (29), or other environmental exposures can alter host interactions with microbes. Differences in microbiome characteristics may also persist via mother-to-child transmission, as infant microbiomes are seeded from the birth canal and/or breastfeeding (30, 31). Further, social network homophily and shared built environments may represent reservoirs of shared microbiota membership (32).

So far, limited research has examined sociodemographic associations with the oral microbiome. The Human Microbiome Project (HMP) collected microbiome samples at nine distinct oral sites on a volunteer sample in the U.S. with minimal race/ethnic variability (approx. 80% white) (33, 34). Nonetheless, the HMP found differentially abundant taxa comparing non-Hispanic white, non-Hispanic black, Asian, Mexican, and Puerto Rican ethnicities (35). In another U.S. volunteer sample, distinct subgingival microbiomes were identified by race/ethnicity, with non-Hispanic blacks having lower microbiome diversity than other groups (36). In a comparison of salivary microbiomes of Cheyenne and Arahapo vs. non-Native individuals in the U.S., strong bacterial species composition clustering, differences in species richness, and numerous differentially abundant taxa were found by ethnicity (37). Several low-throughput studies examining specific periodontal pathogens found significant differences in abundance and/or presence by race/ethnicity (38-40). To our knowledge, only one study has tested associations between SES and the oral microbiome, finding substantial differences (20% of variation) by municipal-level SES in the Danish Health Examination Survey (41).

In order to explore the relationship between the oral microbiome and health disparities, population-level sociodemographic associations must be assessed. Our aim was to assess sociodemographic variation in the human salivary microbiome. Specifically, we examined whether bacterial taxa were differentially abundant, and whether variation existed in alpha and beta diversity by sociodemographic characteristics using high-throughput sequencing data from a population-based sample.

## 3 Methods

### 3.1 Data Source

Samples came from the 2013-14 New York City Health and Nutrition Examination Survey (NYC HANES-II) previously described (42). Briefly, the 2013-14 NYC HANES was the second population-representative, cross-sectional survey of adult NYC residents, using a three-stage cluster sampling design. Overall response rate was 36% (n=1524). Eligible participants completed a two-part interview, physical examination, and blood, urine, and oral mouthwash biospecimen collection. Nearly all participants (95%) provided an oral mouthwash specimen. This study was approved by the institutional review boards of the City University of New York and the New York City Department of Health and Mental Hygiene, and all participants gave informed consent. Participants providing mouthwash specimens in the current sub-study also consented to use these specimens in future studies.

### 3.2 Subsample Selection

The current study uses 297 NYC HANES participants selected to examine oral microbiome associations with tobacco use, as described elsewhere [CITATION PENDING – Beghini 2018 Companion Paper]. Briefly, we selected the 90 self-reported current cigarette smokers with the highest serum cotinine, 45 randomly selected never smokers with serum cotinine <0.05 ng/mL, 45 randomly selected former smokers with serum cotinine <0.05 ng/mL, all 38 former and never smokers with serum cotinine between 1 and 14 ng/mL, and 79 participants reporting usage of hookah, cigar, cigarillo and/or e-cigarette in the last 5 days. Descriptive statistics in the subsample and overall NYC HANES sample are presented in Table 1.

**Table 1.** Demographics

|  | Oral Microbiome Subsample | Full NYC HANES Sample |
| --- | --- | --- |
| Total | 282 | 1527 |
| Age in years – median [range] | 42 [20 to 94] | 42 [20 to 97] |
| Age group (%) |  |  |
| 20-29 | 70 (24.8) | 360 (23.6) |
| 30-39 | 60 (21.3) | 337 (22.1) |
| 40-49 | 51 (18.1) | 252 (16.5) |
| 50-59 | 51 (18.1) | 264 (17.3) |
| 60 and over | 50 (17.7) | 314 (20.6) |
| Sex = Female (%) | 150 (53.2) | 885 (58.0) |
| Educational achievement (%) |  |  |
| College graduate or more | 87 (30.9) | 628 (41.1) |
| Less than High school diploma | 65 (23.0) | 316 (20.7) |
| High school graduate/GED | 63 (22.3) | 244 (16.0) |
| Some College or associate's degree | 67 (23.8) | 337 (22.1) |
| Missing | 0 ( 0.0) | 2 ( 0.1) |
| Annual family income (%) |  |  |
| \$60,000 or more | 82 (29.1) | 429 (28.1) |
| Less Than \$30,000 | 105 (37.2) | 537 (35.2) |
| \$30,000 - \$60,000 | 59 (20.9) | 348 (22.8) |
| Missing | 36 (12.8) | 213 (13.9) |
| Marital Status (%) |  |  |
| Married | 96 (34.0) | 590 (38.6) |
| Widowed | 15 ( 5.3) | 76 ( 5.0) |
| Divorced | 23 ( 8.2) | 156 (10.2) |
| Separated | 12 ( 4.3) | 51 ( 3.3) |
| Never married | 101 (35.8) | 511 (33.5) |
| Living with partner | 35 (12.4) | 143 ( 9.4) |
| Race/ethnicity (%) |  |  |
| Non-Hispanic White | 97 (34.4) | 513 (33.6) |
| Non-Hispanic Black | 75 (26.6) | 340 (22.3) |
| Hispanic | 71 (25.2) | 390 (25.5) |
| Asian | 22 ( 7.8) | 204 (13.4) |
| Other | 17 ( 6.0) | 80 ( 5.2) |
| Place of birth (%) |  |  |
| US, PR and Territories | 90 (31.9) | 668 (43.7) |
| Other | 190 (67.4) | 851 (55.7) |
| Missing | 2 ( 0.7) | 8 ( 0.5) |
| Gum disease (self-reported) (%) |  |  |
| Yes | 27 ( 9.6) | 175 (11.5) |
| No | 254 (90.1) | 1322 (86.6) |
| Missing | 1 ( 0.4) | 30 ( 2.0) |
| Mouthwash use (times per week) (%) |  |  |
| None | 115 (40.8) | 591 (38.7) |
| 1 to 5 | 68 (24.1) | 370 (24.2) |
| 6 to 7 | 99 (35.1) | 565 (37.0) |
| Missing | 0 ( 0.0) | 1 ( 0.1) |
| Sugar-sweetened beverages (per week) (%) |  |  |
| 0-<1 | 152 (53.9) | 985 (64.5) |
| 1-5 | 67 (23.8) | 313 (20.5) |
| 6 or more | 62 (22.0) | 227 (14.9) |
| Missing | 1 ( 0.4) | 2 ( 0.1) |
| Smoking status (%) |  |  |
| Cigarette | 86 (30.5) | 215 (14.1) |
| Never smoker | 43 (15.2) | 843 (55.2) |
| Former smoker | 43 (15.2) | 285 (18.7) |
| Alternative smoker | 72 (25.5) | 142 ( 9.3) |
| Secondhand | 38 (13.5) | 42 ( 2.8) |

### 3.3 Oral rinse collection and microbiome sample processing

Participants were asked to fast for 9 hours prior to oral rinse collection. A 20-second oral rinse was divided into two 5-second swish and 5-second gargle sessions using 15 mLs of Scope^®^ mouthwash. After each session, participants expectorated the rinse into a sterile cup. Timers built into the computer-assisted personal interview program signaled the timing of the swish, gargle and expectoration. Oral rinse specimens were stored cold before delivery to the New York Public Health Laboratory where they were transferred into 50 mL centrifuge tubes, frozen and stored at −80°C. The oral rinse samples were then transported on dry ice to Albert Einstein College of Medicine, where they were stored at −80°C until processing.

Specimen processing and sequence analysis methods are described in detail in the appendix. Briefly, we extracted DNA using QIAamp DNA mini kit (QIAGEN), and amplified DNA in the V4 region of the 16S rRNA using primers 16SV4_515F (GTGYCAGCMGCCGCGGTA) and 16SV4_806R (GGACTACHVGGGTWTCTAAT) (38,39), followed by amplicon sequencing using a MiSeq (Illumina, San Diego, CA) with 2×300 paired-end fragments. We analyzed 16S reads using QIIME version 1.9.1 (40) and Phyloseq (41). We merged raw Illumina paired-end reads using the QIIME command fastq-join (42), and discarded any resulting low quality reads (PHRED score < 30) when joining the split reads (qiime split_libraries_fastq.py). We performed open-reference Operational Taxonomic Unit (OTU) picking by clustering using UCLUST at 97% sequence similarity, and we assigned taxonomy using the SILVA 123 (43) database. We removed samples with less than 1000 reads (n=15) from the OTU table and collapsed genera present with a mean relative abundance of less than 2 □ 10-4 into a category labelled “Other.” (43-46)

### 3.4 Statistical Analysis

We compared differences in oral microbiome characteristics by seven sociodemographic factors (race/ethnicity, age, group, sex, educational attainment, income tertiles, marital status, nativity) and by several behavioral/oral health measures: diet (sugar sweetened beverages, meat, poultry, fish, vegetables, and fruits, recorded as times consumed in the past week); oral health behaviors (mouthwash use, flossing, time since last dental visit) and smoking status (categories defined above). We assessed pairwise correlation between sociodemographic variables using Cramer’s V, a correlation coefficient for nominal variables.

To assess differential abundance by sociodemographic variables, we used edgeR (47) to estimate a series of log-linear generalized linear models (GLMs) predicting each OTU abundance. OTUs were considered differentially abundant at false discovery rate (FDR) < 0.01.

Before edgeR, we filtered out OTUs that did not have three or more samples with a count of at least eight, leaving 216 OTUs for analysis, a filter representing an approximate inflection point on the curve of remaining OTUs against the minimum count. To examine potential mediators, we fit crude models as well as models adjusted for oral health behaviors, diet, smoking status, and age and sex (when applicable). edgeR was conducted at the taxonomic level of highest specificity allowed, which was the genus in all cases where FDR was less than 1%; therefore differential abundance findings are presented at the genus level.

We measured alpha diversity using Chao1 richness (48), which we compared by each sociodemographic variable using Kruskal-Wallis tests. Beta diversity was assessed using principal coordinates analysis and permutation multivariate analysis of variance (PERMANOVA) (49) on weighted UniFrac distances (50). To ensure results were not driven by selection on smoking status, we also compared alpha and beta diversity adjusting for smoking status.

We performed clustering of samples with respect to OTUs using partitioning around medoids on Bray Curtis, Jenson-Shannon, root-Jenson Shannon, weighted and unweighted UniFrac distances (51). Prediction strength (PS) was calculated for k=2:10 clusters on each distance measure, using PS≥0.9 to signify strong support for k clusters (51).

Statistical analyses were conducted in R version 3.4 (52) for Linux.

## 4 Results

### 4.1 Descriptive Statistics

The initial subsample included 297 participants; after removing samples with less than 1000 reads, there were 282 participants remaining for analysis. Table 1 shows descriptive statistics for sociodemographic characteristics including age (median [range]: 42 [20 to 94]), sex (53.2% female), race/ethnicity (34.4% non-Hispanic White, 26.6% non-Hispanic Black, 25.2% Hispanic), annual family income (42.7% less than $30K, 33.3% $60k or more), and educational achievement (23.0% less than high school diploma, 30.9% college degree or greater). Cramer’s V on all pairwise combinations of sociodemographic variables showed only minor collinearity (all V<.35) (Figure A1), indicating associations with the microbiome for each sociodemographic variable do not merely reflect correlations between sociodemographic variables.

### 4.2 Relative Abundance and Alpha Diversity

Oral microbiomes were characterized at the phylum level by a gradient between Firmicutes and Bacteroides abundance, with overall dominance by Firmicutes (mean=52±10%). Streptococcus was the most abundant genus (36±10%) followed by Prevotella (17±8%). (Figure 1). The overall mean Chao1 was 462±118, with no differences by age group (p=0.79), sex (p=0.13), educational achievement (p=0.92), annual family income (p=0.62), marital status (p=0.54), race/ethnicity (p=0.13), or nativity (p=0.97) (Figure A2). These results were not changed by adjustment for smoking.

**Figure 1.**
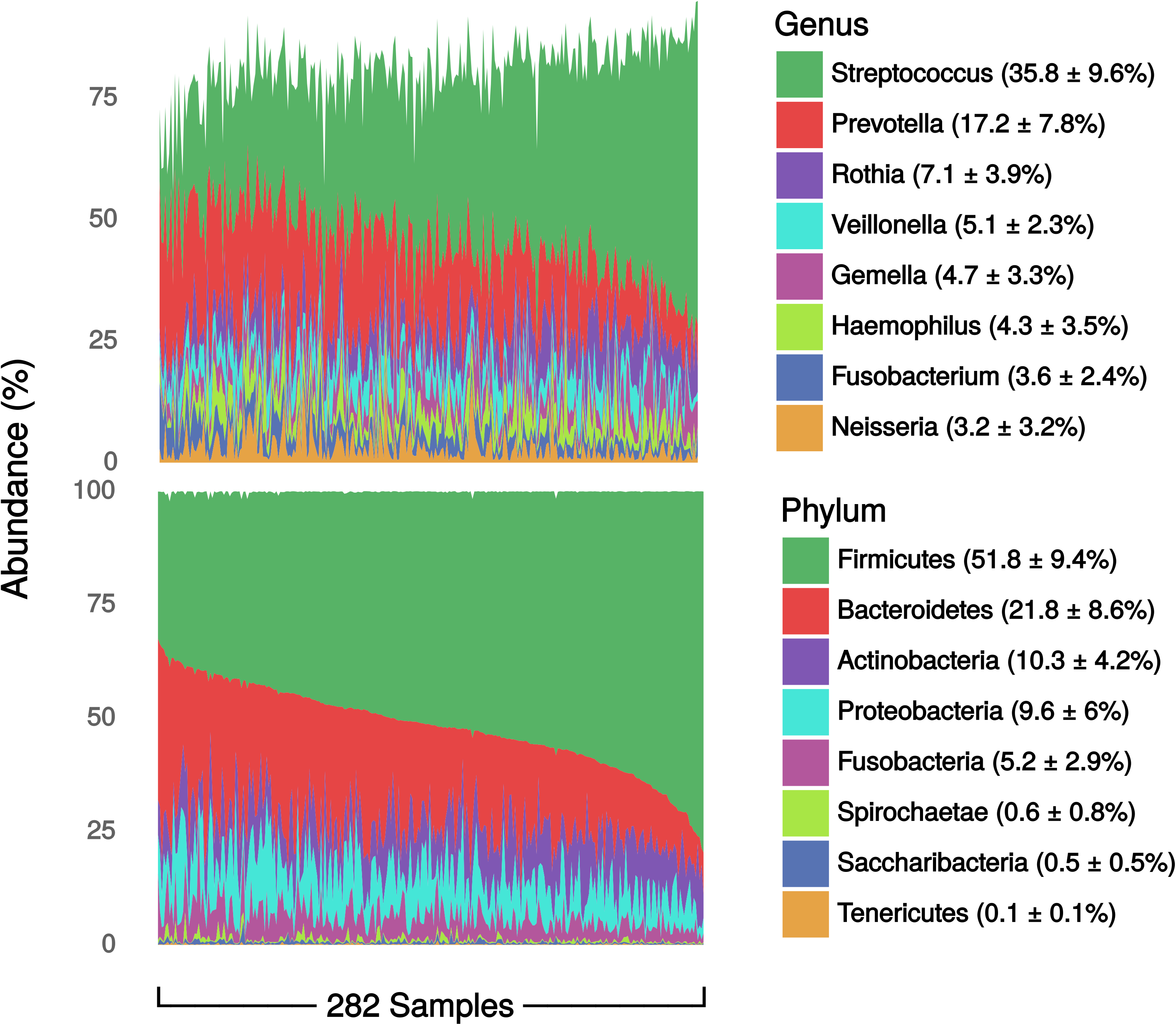
Genus- and phylum-level relative abundances. Data are percent of overall communities within samples, summarized as mean ± standard deviation of percent across samples. Data are from the oral microbiome subsample (n=282) of the New York City Health and Nutrition Examination Survey, 2013-2014.

### 4.3 Differential Abundance

Numerous taxa were differentially abundant by race/ethnicity, nativity, marital status, gender, family income, education, and age. Figure 2 displays log-base-2 fold change (logFC), or coefficient from edgeR log linear models, for each comparison group and all significant OTUs. The logFC can be interpreted as the log-base-2 ratio of relative abundance compared to the reference group, so that e.g. *Lactobacillus* is found to be 2^2.5^ = 5.7 times as abundant among participants with family incomes of $30-60,000 per year, compared to $60,000 or more. A total of 69 OTUs were differentially abundant by any sociodemographic variable, including 56 by age group, 27 by race/ethnicity, 21 by family income, 19 by education, 19 by marital status, seven by nativity, and three by sex. We also found 12 unique OTUs differentially abundant by oral health behaviors, 49 by smoking status, and 23 by diet variables. The most frequently differentially abundant were Lactobacillus (all variables), and Prevotella (age, education, family income, marital status, race/ethnicity, nativity, Figure 2). Differential abundance findings for selected taxa are presented in Table 2 (see table A1 for all differential abundance findings).

**Figure 2.**
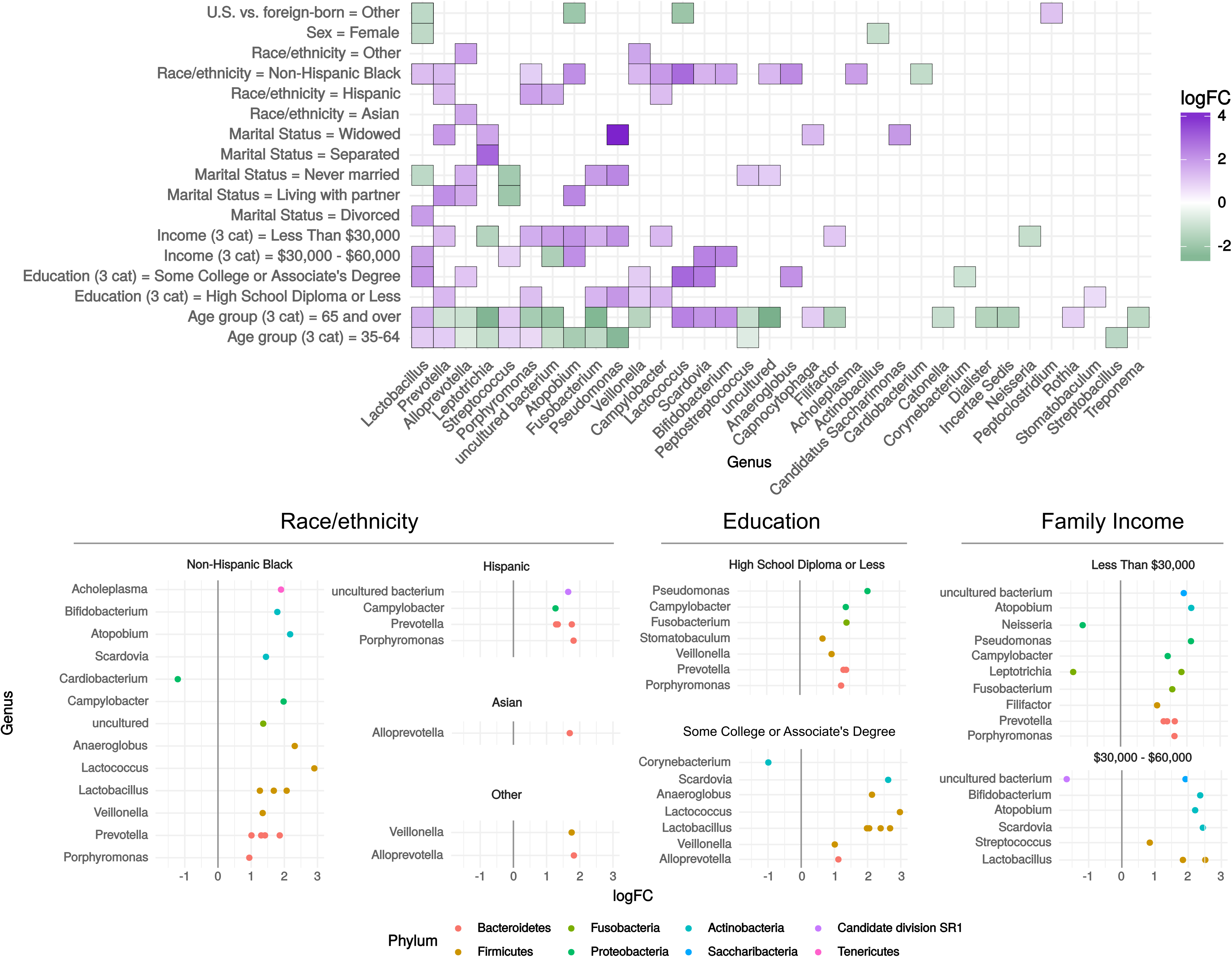
Differential abundance by sociodemographic characteristics. OTUs meeting unadjusted FDR < 0.01 in negative binomial log-linear GLMs using edgeR. Data are from the oral microbiome subsample (n=282) of the New York City Health and Nutrition Examination Survey, 2013-2014. Filled tiles in (A) indicate the genus had at least one OTU differentially abundant by at least one coefficient contrast within the sociodemographic factor. Where more than one OTU was significant within one genus, the maximum logFC is displayed in (A). Reference groups for sociodemographic variables are as follows: Sex: Male, Age: 20-34, Education: College Graduate or More, Family income: $60,000 or more, Marital status: Married, Race/ethnicity: Non-Hispanic White, US-vs. foreign-born: US-Born, 50 States, DC, PR and Territories. Abbreviations: cat=categories; GLM=generalized linear model; logFC=log fold change; OTU=operational taxonomic unit; US=United States.

**Table 2.** Differential abundance findings for OTUs selected based on clinical relevance, where FDR < 0.01. Data are from the oral microbiome subsample (n=282) of the New York City Health and Nutrition Examination Survey, 2013-2014.

|  | **Lactobacillus** |  | **Prevotella** |  | **Streptococcus** |  | **Porphyromonas** |  | **Fusobacterium** |  | **Lactococcus** |  |
| --- | --- | --- | --- | --- | --- | --- | --- | --- | --- | --- | --- | --- |
|  | **logFC** | **FDR** | **logFC** | **FDR** | **logFC** | **FDR** | **logFC** | **FDR** | **logFC** | **FDR** | **logFC** | **FDR** |
| **Race/ethnicity** |  |  |  |  |  |  |  |  |  |  |  |  |
| Non-Hispanic White | 0 | *Ref* | 0 | *Ref* | 0 | *Ref* | 0 | *Ref* | 0 | *Ref* | 0 | *Ref* |
| Non-Hispanic Black | 2.1 | <0.0001 | 1.9 | <0.0001 | - | - | 0.9 | 0.0018 | - | - | 2.9 | <0.0001 |
| Hispanic | - | - | 1.8 | <0.0001 | - | - | 1.8 | <0.0001 | - | - | - | - |
| **Family income** |  |  |  |  |  |  |  |  |  |  |  |  |
| \$60,000 or more | 0 | *Ref* | 0 | *Ref* | 0 | *Ref* | 0 | *Ref* | 0 | *Ref* | 0 | *Ref* |
| \$30,000 - \$60,000 | 2.5 | <0.0001 | - | - | 0.9 | 0.003 | - | - | - | - | - | - |
| Less Than \$30,000 | - | - | 1.6 | 0.0025 | - | - | 1.6 | <0.0001 | 1.5 | 0.003 | - | - |
| **Education** |  |  |  |  |  |  |  |  |  |  |  |  |
| College Graduate or More | 0 | *Ref* | 0 | *Ref* | 0 | *Ref* | 0 | *Ref* | 0 | *Ref* | 0 | *Ref* |
| Some College or Associate's Degree | 2.7 | <0.0001 | - | - | - | - | - | - | - | - | 3.0 | <0.0001 |
| High School Diploma or Less | - | - | 1.4 | 0.0064 | - | - | 1.2 | 0.0008 | 1.4 | 0.006 | - | - |

Abbreviations: logFC, log fold change; FDR, false discovery rate; Ref, reference group. “-“ indicates genus was not significantly differentially abundant.

Figure 3 displays the boxplots of absolute values of logFCs for both crude and adjusted models. The OTUs selected for display in all models are the OTUs meeting FDR <0.01 in crude models. Comparing adjusted vs. crude boxplots allows a visual assessment of the effect of adjustment on the entire set of OTUs: a shift towards zero reflects attenuation while a shift away from zero reflects amplification. Over all sociodemographic variables, a minor attenuating effect was observed after adjusting for smoking (mean change in logFC, −3.9%), oral health behaviors (−4.9%), diet (−6.3%), age and sex (−3.3%). Adjustment for oral health had the largest impact on logFCs for age group (−4%), sex (−27.4%), and nativity (−13.5%); diet had the strongest impact on logFCs for education (−13.1%) and marital status (−16.9%), smoking had the strongest impact on logFCs for family income (−11.9%), and age and sex had the strongest impact on logFCs for race/ethnicity (−4.2%).

**Figure 3.**
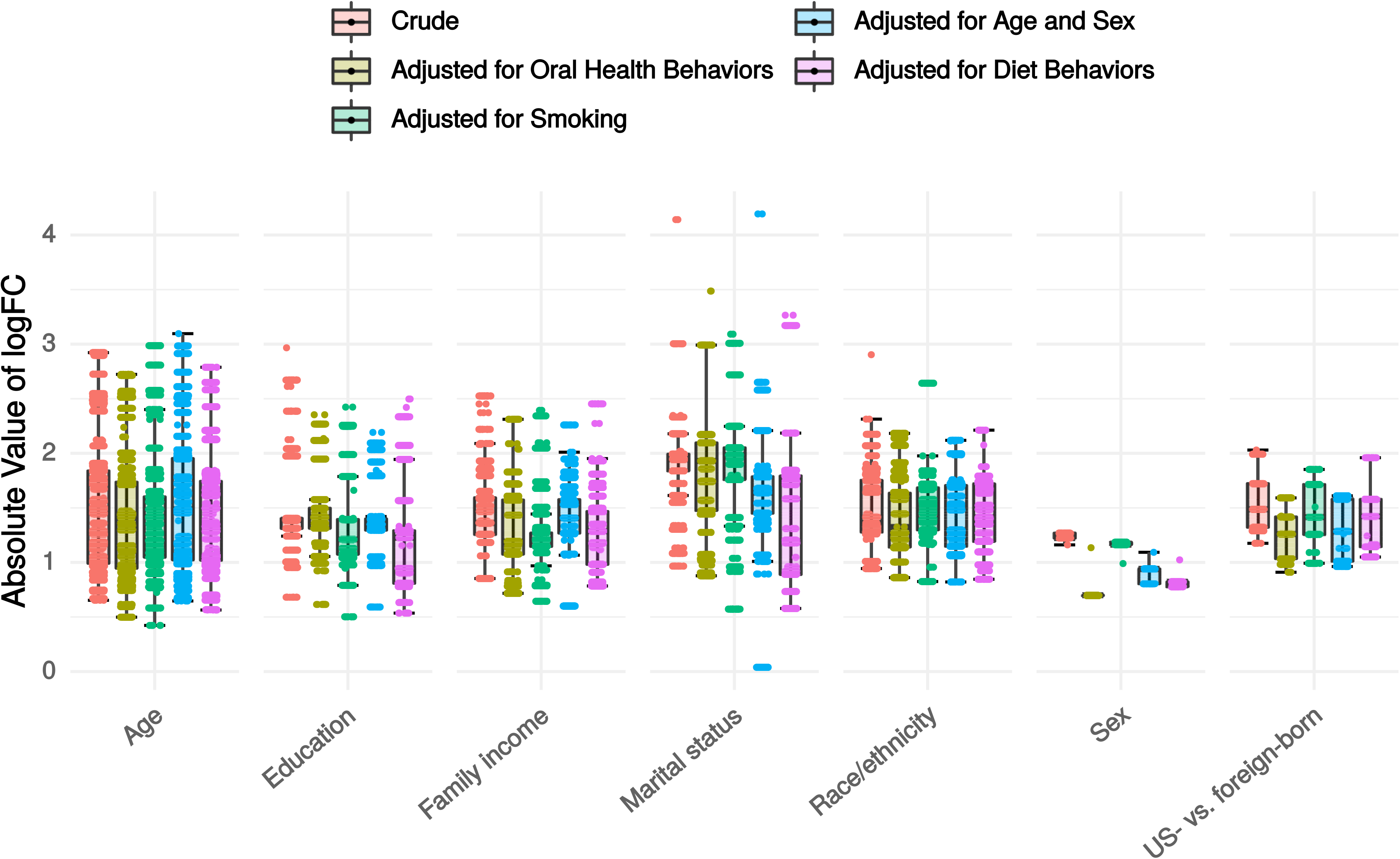
Distribution of absolute values of log-fold change (logFC) in crude and adjusted negative binomial log-linear GLMs edgeR models for each sociodemographic variable. Data are from the oral microbiome subsample (n=282) of the New York City Health and Nutrition Examination Survey, 2013-2014. Abbreviations: GLM=generalized linear model; logFC=log fold change; US=United States.

### 4.4 Beta Diversity and Clustering

Figure 4 illustrates between-versus within-group weighted UniFrac distances by each sociodemographic variable. We observed overall shifts in composition by age group (p=0.017, r^2^=0.026), with no other variables showing greater between-than within-group variation, a result which was not changed by adjusting for smoking. Plots of the first two principal coordinates based on weighted UniFrac distances showed little patterning by any variable (not shown). Clustering scores were sensitive to the distance metric used, with Bray-Curtis indicating moderate support for 2 clusters (PS=0.86), and all other measures providing little support for clustering.

**Figure 4.**
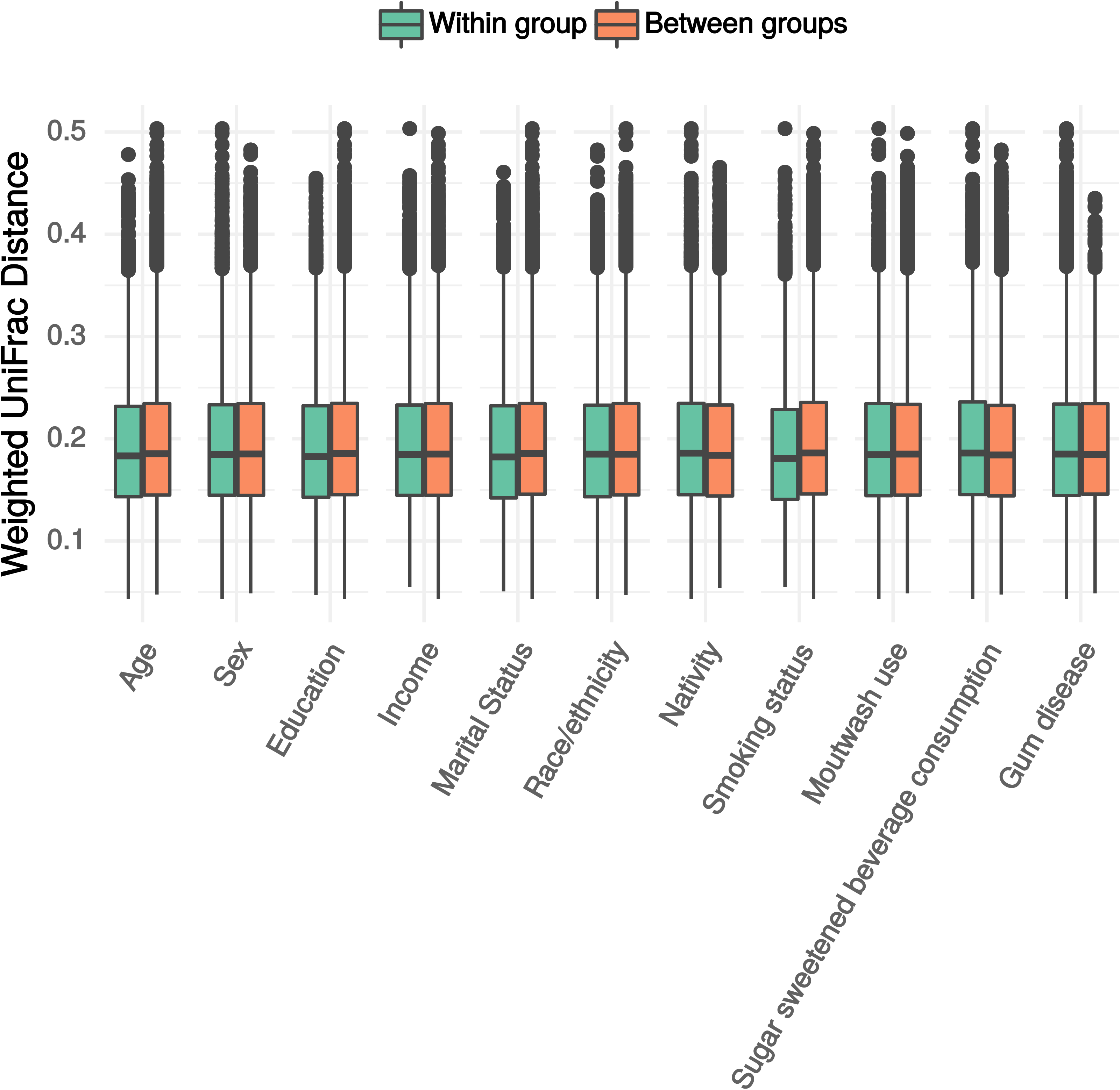
Within and between group beta diversity estimate distributions. Data are from the oral microbiome subsample (n=282) of the New York City Health and Nutrition Examination Survey, 2013-2014. Abbreviations: cat=category.

## 5 Discussion

In a diverse population-based sample, we found that a large number of bacterial taxa in the oral microbiome were differentially abundant by age, race/ethnicity, family income, education, nativity, and sex. Notably, we found a greater number of associations with SES variables (21 by family income, 19 by education) than with sex, marital status or nativity. There were also more associations with SES than oral health behaviors (12). Sociodemographic associations were not appreciably diminished by adjustment for smoking, oral health behaviors, or dietary behaviors. Alpha diversity was similar across groups, and beta diversity explained only a small proportion of variance by age (2.7%), and less by other variables.

Many genera found differentially abundant by multiple variables represent taxa that have documented associations with health and disease. *Streptococcus, Lactobacillus* (53),*Prevotella* (54) *Fusobacterium* (55), and *Porphyromonas* (56, 57) are understood to play a role in oral disease. Further, many of these organisms likely play a role in wide ranging systemic conditions (15). Specifically, *Fusobacterium spp.* have been linked to colorectal cancer (58, 59), adverse pregnancy outcomes, CVD and rheumatoid arthritis (60). *Porphyromonas gingivalis* is a key determinant of oral microbiome structure (61), and is hypothesized to mediate an array of systemic pathogenic processes (15), including associations with stroke (11), CHD (12), a number of cancers (17, 18, 62) and rheumatoid arthritis (22).

To our knowledge, our study is the first to examine differences in the oral microbiome by individual level sociodemographic factors in a population-based sample. Our finding of differentially abundant taxa by race/ethnicity is consistent with previous studies with small volunteer samples. The HMP Consortium found that, for all body sites, ethnicity was the host phenotypic variable with the most associations (35). For the oral microbiome, a study examining 40 periodontal disease-related taxa found differences among Asian, Hispanic, and blacks (38). Two lower-throughput studies found greater *Prevotella* and *Porphyromonas* prevalence (40), and lower *Fusobacterium* abundance (39) in blacks vs. whites. Our finding of differential OTUs by SES variables is also consistent with findings from the Danish Health Examination Survey (DANHES, n=292), which found nine differentially abundant taxa by municipal-level SES (41).

Adjustment for smoking, diet, and oral health behaviors each exerted a moderate attenuating effect on differential abundance findings across sociodemographic categories. This stands to reason in light of findings by our group [CITATION PENDING – Beghini 2018 Companion Paper] and others (29) that smoking is associated with major shifts in the oral microbiome, along with similar findings for diet (63), and indicates that some portion of observed sociodemographic patterning reflects differences in health habits or access to dental care. However, the finding that differential abundance was not eliminated by adjustments suggests that additional mechanism underlie sociodemographic variation in the oral microbiome. These may include upstream social factors such as psychosocial stress (27) or features of the built environment (32).

While existing oral microbiome studies are limited, the absence of differences in alpha and beta diversity by race/ethnicity contrasts with two previous studies among non-population-based samples. These found differences in alpha diversity and ethnicity-based clustering in oral microbiomes in non-Hispanic Blacks vs. Whites (36), and in Cheyenne and Arahapo vs. non-native individuals (37). Differences in alpha and beta diversity can indicate larger-scale shifts in composition; our finding that specific OTUs were differentially abundant but that overall shifts were less present may indicate that, at a population level, sociodemographic patterns in oral microbiome composition are more subtle.

### 5.1 Limitations

Despite the strength of NYC-HANES as a diverse population-based sample, the cross-sectional design limits its ability to test the oral microbiome as a mediator in health disparities, as changes in the oral microbiome may reflect existing disease rather than etiological factors. Additionally, our findings are limited by having primarily genus-level information, and in many cases salient differences exist at a greater degree of taxonomic specificity – for example, with *P. gingivalis*, *F. nucleatum*, and *Prevotella intermedia*. There may also be wide variability in virulence even at the species level, as is the case with *P. gingivalis* (64). Given the importance of many of the differentially abundant genera in health and disease, our findings suggest that further investigation into the role of the oral microbiome in health disparities is warranted. Future investigations should consider use of whole genome shotgun sequencing or other methods able to provide more specific taxonomic classification and describe functional, as well as taxonomic, composition.

### 5.2 Conclusion

Our results lend support to potential role of the social environment in shaping microbiome composition at the population level (24, 65). The finding of differentially abundant OTUs, many of which are health-relevant, for every sociodemographic variable, suggests that these associations may be important in determining population health patterns. In particular for race and SES, but also for nativity and marital status, the finding that multiple health-relevant microbes are differentially abundant supports a growing hypothesis that the microbiota may partially mediate long-observed social disparities in major disease outcomes. At a minimum, these results highlight that social factors may be important potential confounders in studies of the human oral microbiome and health.

Mechanisms for the observed associations are currently unknown, and one important next step will be to examine the multiple levels of exposures underlying these associations, including macro-level social and health policy, exposure to psychosocial stressors, outdoor and built environment features, and social interactions (24). Importantly, if the microbiome is a partial mediator of health disparities, then identifying modifiable features of the social environment that are most strongly associated with the microbiome can inform effective interventions to improve population health and reduce health disparities.

## 8 Acknowledgements

The individual author contributions are as follows: HEJ, LW, LT, RB, and JD conceptualized and designed the study; AR, FB, NS, and LW led data analysis and data visualization; RB, CPZ, MU, and TUM led specimen processing and 16S data generation; AR wrote first draft of manuscript; and all authors contributed to editing/revisions on manuscript. We gratefully acknowledge the efforts of the New York City Department of Health and Mental Hygiene in co-leading the parent NYC HANES study. In particular, we wish to thank Sharon Perlman, Carolyn Greene, Claudia Chernov, Amado Punsalang, and the many other staff who helped support data collection.

---

Conflict of Interest and Authorship Conformation Form

Please check the following as appropriate:

⊠ All authors have participated in (a) conception and design, or analysis and interpretation of the data; (b) drafting the article or revising it critically for important intellectual content; and (c) approval of the final version.
⊠ This manuscript has not been submitted to, nor is under review at, another journal or other publishing venue.
⊠ The authors have no affiliation with any organization with a direct or indirect financial interest in the subject matter discussed in the manuscript
□ The following authors have affiliations with organizations with direct or indirect financial interest in the subject matter discussed in the manuscript:

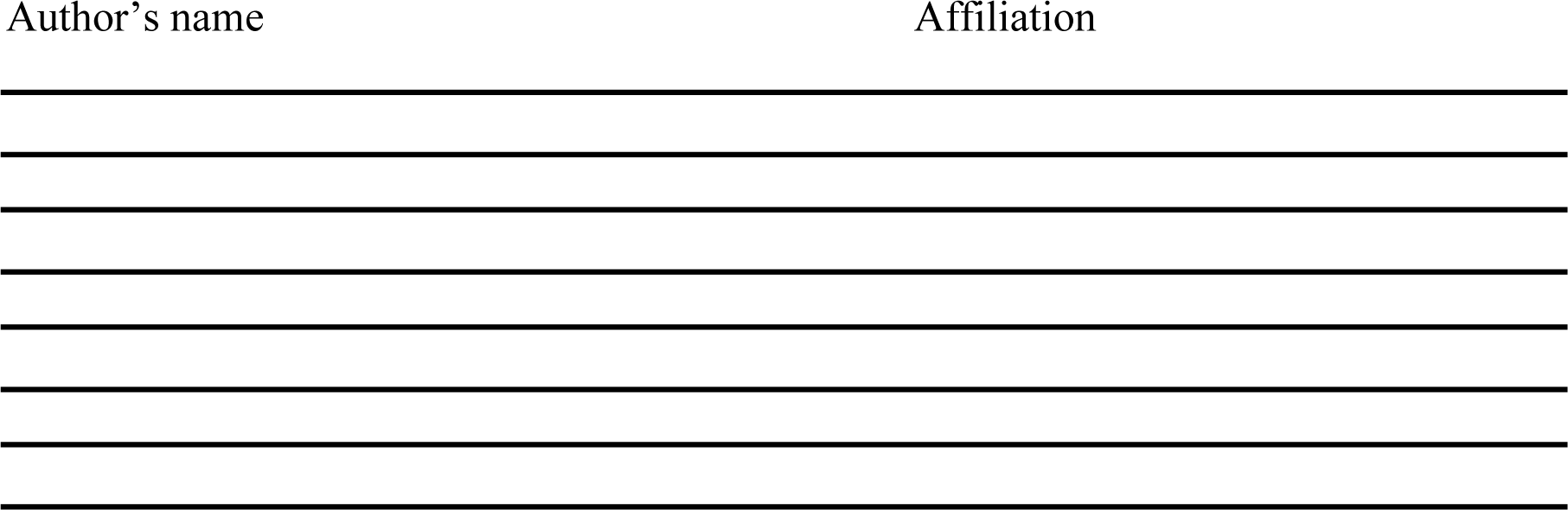

## DNA Extraction

All laboratory procedures were performed under a hood (AirClean Systems) to minimize environmental contamination and negative controls were used throughout. From each oral rinse sample, a 1.5 mL aliquot was centrifuged at 750 x g for 5 min and all but 150 μl of supernatant was removed. The pellet was re-suspended in the remaining supernatant and incubated in an enzyme mixture consisting of lysozyme (0.84 mg/ml, Sigma Aldrich), mutanolysin (0.25 U/ml, Sigma Aldrich) and lysostaphin (21.10 U/ml, Sigma Aldrich), at 37°C for 30 minutes. This was followed by incubation at 56°C for 10 minutes in 15 μl proteinase K and 150 μl Buffer AL. Samples were then transferred to screw top tubes with 100 g of 0.1-mm-diameter Zirconia/Silica Beads (BioSpec) and bead beaten using a FastPrep-24 homogenizer (MP Biomedicals) at speed 6.0 for 40 seconds. Tubes were centrifuged at 750 x g for 30 sec and 150 μl of supernatant was added to a new 1.7 ml tube with 150 μl of 100% ethanol and mixed by vortexing for 15 seconds. Supernatant was then added to the spin column from the QIAamp DNA mini kit (QIAGEN) and centrifuged at 6000 x g for 1 minute. Column purification was performed according to the QIAamp DNA mini kit directions starting at the AWI wash step. Final elution was performed in 100 μl of Buffer AE.

## 16S rRNA Gene Amplification

DNA was amplified for the V4 variable region of the 16S rRNA gene using the primers 16SV4_515F (GTGYCAGCMGCCGCGGTA) and 16SV4_806R (GGACTACHVGGGTWTCTAAT) (45, 46). Each primer had an 8-bp unique Hamming barcode with forward primers containing a 3-bp (TCG) and 4-bp (ACTG) pad on either side, with reverse primers including a 3-bp (GTA) and 4-bp (TC) pad on each side of the barcode (47). PCR reactions were performed with 17.75 μl of nuclease-free PCR-grade water, 2.5 μl of 10X Buffer w/ MgCl2 (Affymetrix, Santa Clara, CA), 1μl of MgCl2 (25 mM, Affymetrix, Santa Clara, California, USA), 0.5 μl of dNTPs (10 mM, Roche, Basel, Switzerland), 0.25 μl of AmpliTaq Gold DNA Polymerase (5 U/μl, Applied Biostystems, Foster City, California), 0.5 μl of HotStart-IT FideliTaq (2.5 U/μl, Affymetrix, Santa Clara, CA), 1μl of each primer (5 μM), and 0.5 μl of DNA extraction template. Thermal cycling conditions consisted of initial denaturation of 95°C for 5 min, followed by 15 cycles of 95°C for 1 min, 55°C for 1 min, and 68°C for 1 min, followed by 15 cycles of 95°C for 1 min, 60°C for 1 min, and 68°C for 1 min, a final extension for 10 min at 68°C on a GeneAmp PCR System 9700 (Applied Biosystems, Foster City, CA).

PCR products were combined before running 100 μl of the pooled products on a 4% agarose gel at 80V for 2 hours. The ~450 bp bands were excised from the gel and purified using a QIAquick Gel Extraction Kit (Qiagen, Hilden, Germany) and eluted in 30 μl of elution buffer. Purified PCR products were quantified using a Qubit 2.0 Fluorometic High Sensitivity dsDNA Assay (Life Technologies, Carlsbad, CA).

## Library Preparation and Sequencing

Library preparation of the purified PCR products was performed using a KAPA LTP Library Preparation Kit (Kapa Biosystems, Wilmington, MA). The size integrity of the amplicon was validated with a 2100 Bioanalyzer (Agilent Technologies, Santa Clara, CA). High-throughput amplicon sequencing was conducted on a MiSeq (Illumina, San Diego, CA) using 2×300 paired-end fragments. The fastq sequences from the Illumina MiSeq were demultiplexed using Novobarcode (Novocrat Technologies, Selangor, Malaysia) and the 5’-pads and primers were trimmed from each read.

Bacterial taxa were determined by clustering the 16S rRNA sequences into operational taxonomical units (OTUs) using 97% similarity, taxonomy was assigned at the genus level using the SILVA 123 (43) database as reference, excluding samples with less than 1000 reads.

---

**Figure A1.**
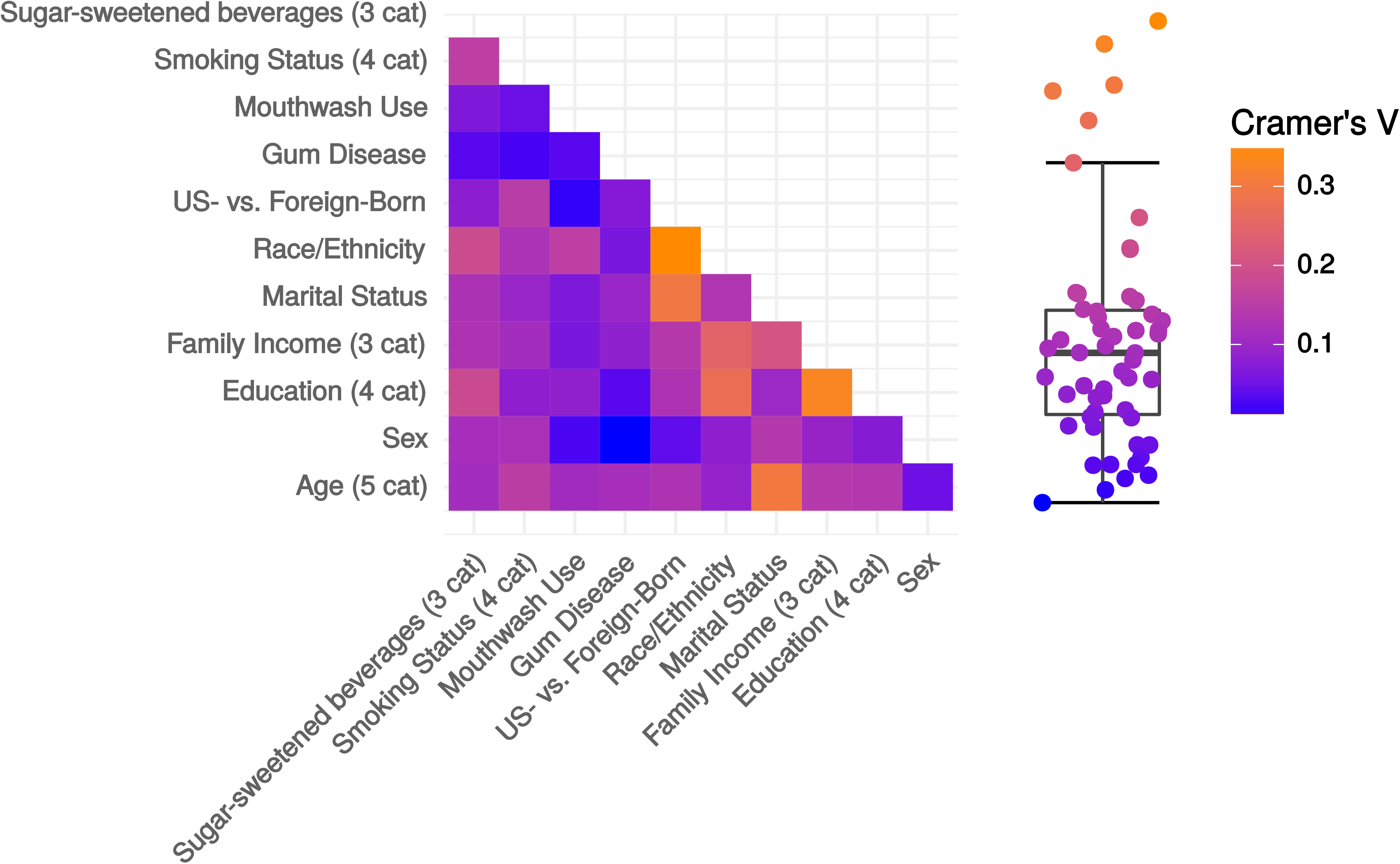
Examining collinearity among sociodemographic variables. Data are absolute value of pairwise Cramer’s V correlation coefficient between sociodemographic factor levels. Data are from the full sample (n=1,527) of the New York City Health and Nutrition Examination Survey, 2013-2014. Abbreviations: cat=categories; US=United States.

**Figure A2.**
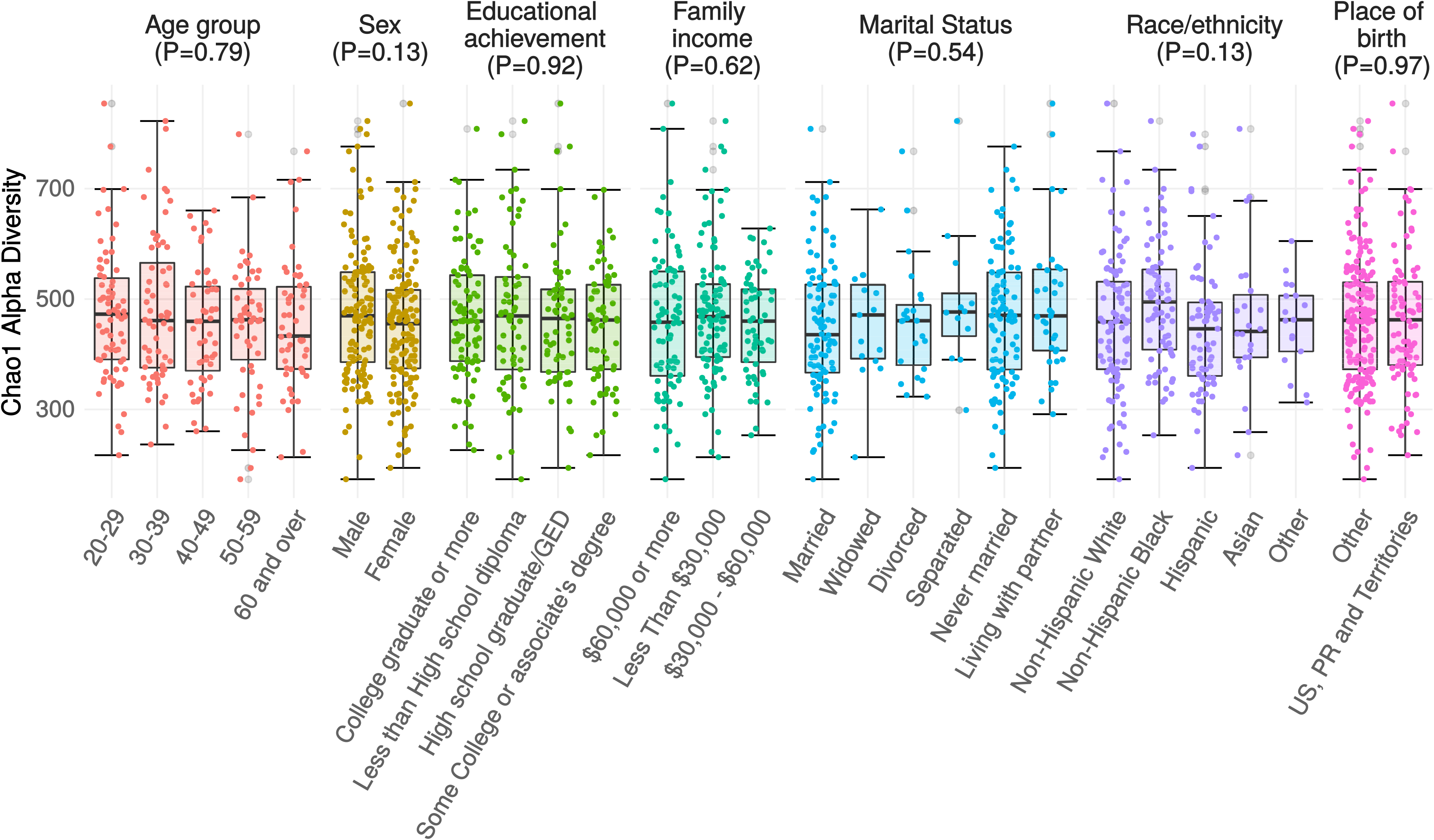
Alpha diversity by Sociodemographic Characteristics. Chao1 alpha diversity of 16S rRNA oral microbiome samples. Measures were compared using a null hypothesis of no difference between groups (Kruskal-Wallis test, p > 0.1 for all tests). Data are from the oral microbiome subsample (n=282) of the New York City Health and Nutrition Examination Survey, 2013-2014. Abbreviations: GED=General equivalency diploma; PR=Puerto Rico; US=United States.

## List of abbreviations

SES: socioeconomic status
CHD: coronary heart disease
CVD: cardiovascular disease
NYC HANES: New York City Health and Nutrition Examination Survey
OTU: operational taxonomic unit
FDR: false discovery rate
PS: prediction strength
logFC: log fold change
HMP: Human Microbiome Project

